# Recording brain activity with ear-EEG (cEEGrids)

**DOI:** 10.1101/2022.10.07.511290

**Authors:** Daniel Hölle, Martin G. Bleichner

**Affiliations:** Neurophysiology of Everyday Life Group, Department of Psychology, University of Oldenburg, Oldenburg, Germany

## Abstract

The cEEGrid (ear-electroencephalography; ear-EEG) is an unobtrusive and comfortable electrode array affixed around the ear. It is suited to investigate brain activity outside of the laboratory for long durations. Previous research established that cEEGrids can be used to study various cognitive processes in and also beyond the lab, even for a whole day. To record high-quality ear-EEG data, careful preparation is necessary. In this protocol, we explain the steps needed for successful experimenting with cEEGrids: First, we show how to test the functionality of the cEEGrid prior to a recording. Second, we describe how to prepare the participant and to fit the cEEGrid, which is the most important step to record high-quality data. Third, we outline how to connect the cEEGrids to the amplifier and how to check the signal quality. In this protocol, we give best practice recommendations and tips that make cEEGrid recordings easier. If researchers follow this protocol, they are comprehensively equipped for experimenting with the cEEGrid in and beyond the lab.

**SUMMARY:** The cEEGrid (ear-electroencephalography) allows to record brain activity in and beyond the lab for extended duration. In this protocol, we describe how to set up and record with cEEGrids.

## INTRODUCTION

With mobile ear-electroencephalography (EEG), brain activity can be recorded in everyday life and new insights into neural processing beyond the lab can be gained ^1^. To be suitable for everyday life, a mobile ear-EEG system should be transparent: unobtrusive, easy to use, motion tolerant, and comfortable to wear even for several hours^2^. The cEEGrid, a c-shaped ear EEG system aims to meet these requirements to minimize interference with natural behavior. The cEEGrid consists of ten Ag/AgCl electrodes printed on flexprint material^3^. Combined with a miniaturized, mobile amplifier and a smartphone for data acquisition^4, 5^, cEEGrids can be used to collect ear-EEG for extended durations^1^.

There are numerous neural processes that can be recorded with electrodes around the ear^6, 7^. Several studies conducted in the lab have shown the potential of the cEEGrid to study these processes. It has been successfully used for auditory attention decoding with above chance level accuracies^8–12^. Segaert and colleagues^13^ used cEEGrids to quantify language impairment in patients with mild cognitive impairment. Garrett and colleagues^14^ showed that cEEGrids are also able to capture auditory brain potentials originating from the brain stem. Apart from research focused on the auditory domain, Knierim and colleagues^15^ used cEEGrids to investigate flow experiences. Pacharra and colleagues^16^ used cEEGrids for a visual Simon Task. All of these lab-based studies showcase the various cognitive processes that can be captured with the cEEGrid.

The cEEGrid can also be used for EEG recordings beyond the lab, as several studies illustrate. For example, cEEGrids have been used for sleep staging^17^, to evaluate mental load in a driving simulator^18, 19^, and to study inattentional deafness, the non-perception of critical alarm sounds, in a flight simulator^20^. In Hölle and colleagues^1^, cEEGrids were used to measure auditory attention during an office day for six hours. In sum, all of these studies highlight the potential of the cEEGrid to investigate various brain processes in and outside of the lab.

Recently, we have complemented our transparent ear-EEG setup with new hardware and software developments. First, we have built the nEEGlace, a mobile EEG amplifier integrated into a neckspeaker^5^. The nEEGlace can be placed around the neck and the cEEGrids can be plugged in directly, making the setup faster and more comfortable for participants. Second, we have developed an Android app, AFEx, that records the current soundscape as privacy-aware acoustic features (signal strength, RMS; spectral properties, PSD; and sound onsets)^21^. These acoustic features can be related to the EEG data and sound processing in everyday life can be studied^21^. Another smartphone app^4^ allows a multitude of different smartphone sensors and other data streams to be recorded concurrently with the EEG signal.

Every EEG recording requires careful preparation to obtain valid results. This is especially important for mobile applications where even more artifacts than in the lab can be expected. To get optimal results with the cEEGrids, specific preparation steps are necessary. We state critical steps in preparing the cEEGrid and the participant for data collection, and how to fit and connect the cEEGrid. We point out potential mistakes and show examples of poor data quality when the cEEGrid is not attached properly. Finally, we show representative results of a piano-played oddball that can be recorded with our Android app.

## PROTOCOL

The general procedure used in this protocol was approved by the ethics board of the University of Oldenburg.

### 1. Testing the cEEGrid

NOTE: If handled with care, cEEGrids can be re-reused several times. To ensure optimal functioning, it should be checked if all electrodes are still working properly before the next recording. We recommend to perform the same procedure for new cEEGrids to spot potential problems with specific electrodes (e.g., due to problems in the manufacturing process) before the recording starts. There are several options to quickly check for problems (e.g. a broken electrode).

1. Option 1: Use a multimeter.
  1.1. Set the multimeter to measure resistance.
  1.2. Attach one pin of the multimeter to the cEEGrid electrode and the other pin to the corresponding contact on the connector end of the cEEGrid.
  1.3. Check if you measure a low resistance (<10kΩ) for each electrode.
2. Option 2: Impedance check of amplifier.
  2.1. Use electrode gel to bridge all cEEGrid electrodes. Make sure that there are no gaps between the electrodes.
  2.2. Attach the cEEGrid to the connector of the amplifier. To see a signal, you need to attach the cEEGrid to the side where you have your reference and ground electrode according to your connector layout.
  2.3. Use the impedance check of your amplifier. Check for impedance of the reference electrode and all eight recording electrodes (10 electrodes total minus ground and reference electrode): they should all have a low impedance (<10kΩ). Afterwards wipe of the gel off the cEEGrid.

### 2. Preparing the participant

NOTE: For high-quality recordings, participant should have clean and dry hair without any hair products (e.g., styling products) and they should not wear make-up. If possible, the participants should wash their hair directly before the recording with a mild and neutral shampoo. Ask your participants to indicate if any of the preparatory steps become uncomfortable for them.

1. Prepare participant
  1.1. Place a cEEGrid around participants’ ear to see how it fits the participant and if it can be positioned around the ear without touching the ear (especially the back of the pinna or the ear lobe, which can be uncomfortable. For larger ears it may be necessary to cut some of the plastic around the electrodes. This pre-fitting also gives an indication of the area that needs to be cleaned.
  1.2. Use abrasive electrode gel to clean the skin around the ear. To gain decent impedance, it is important to clean the skin with some pressure, but make sure it remains comfortable for the participant.
  1.3. Clean this area with alcohol.
  1.4. Dry it off with a clean towel.
  1.5. For higher levels of comfort especially in longer recordings (> 2h), a small piece of tape can optionally be placed on the backside of the ear.
2. Prepare cEEGrid NOTE: There are different ways of attaching the cEEGrid using double-sided tape. We present two options: c-shaped stickers (provided by the manufacturer) that cover the whole surface and small circular stickers that are placed individually around the electrodes (e.g., when reusing cEEGrids).
  2.1. Attach double-sided adhesive stickers (either the c-shaped or individual stickers). Make sure that the stickers do not cover the conductive surface of the electrodes.
  2.2. Put small drops (lentil-sized) of electrode gel on each electrode. Avoid using too much gel, as this might spill over the adhesive material and reduce adhesion to the skin or could create bridges between electrodes.
  2.3. Remove the cover of the adhesive sticker(s). Re-apply gel in case it was removed during this step. Alternatively, you can remove the first cover and apply the gel then; however, this requires a very steady hand so that you do not spill gel accidently on the adhesive)
3. Fit cEEGrid on participant
  3.1. Ask the participant to hold their hair away from the ear so that it does not obstruct the fitting. For longer hair, hair clips can be used. Any hair should be moved out of the way as much as possible (depending on the hairline this is not always possible for the hair directly above the ear), so that the stickers touch the skin directly.
  3.2. Fit the cEEGrid around the ear. Make sure to not place the cEEGrid to close to the ear, as this may become uncomfortable for the participant. Once you have positioned the cEEGrid, press it onto the skin. You may also ask the participant to press onto the electrodes.
  3.3. Repeat for the other ear NOTE: It is important to know the layout of the connector you are using (i.e., position of the ground and the reference electrode). Depending on the system that you are using, the layout may differ.
4. Connect the cEEGrid
  4.1. Connect the cEEGrid connector to the amplifier.
  4.2. Plug the cEEGrid contacts into the connector. Make sure that the contacts are plugged in on the correct side.
5. Check impedance and data.
  5.1. Connect the amplifier to the smartphone (optionally: laptop) via Bluetooth.
  5.2. Check impedance of electrodes. Impedance usually improves over time (5 to 10 minutes) and does not have to be below 10kΩ for each electrode in the beginning. We do not recommend to try to put more gel underneath electrodes with a high impedance. If the impedance of most electrodes and the resulting EEG signal is high, we recommend to fit the entire cEEGrid again.
  5.3. Check the EEG signal. Ask participant to clench their teeth, to blink, and to close their eyes (alpha activity). Observe the corresponding artifacts and alpha activity in the signal. Make sure every electrode provides a good signal.
  5.4. Begin recording.

### 3. Removing cEEGrids and cleaning up

1. Once the recording is completed, disconnect the phone (or laptop) from the amplifier. Detach the cEEGrids from the amplifier and remove the amplifier from the participant. Gently remove the cEEGrids from the participant. Participants can clean themselves with tissues or a towel.
2. Soak cEEGrids in water for some minutes. They can be submerged completely.
3. Carefully detach stickers from the cEEGrid to avoid damage. Rinse off any remaining gel. Dry cEEGrids.

### REPRESENTATIVE RESULTS

1. impedance.svg
2. data_example.svg
3. cEEGrid_oddball.svg

## FIGURE AND TABLE LEGENDS

**1.**
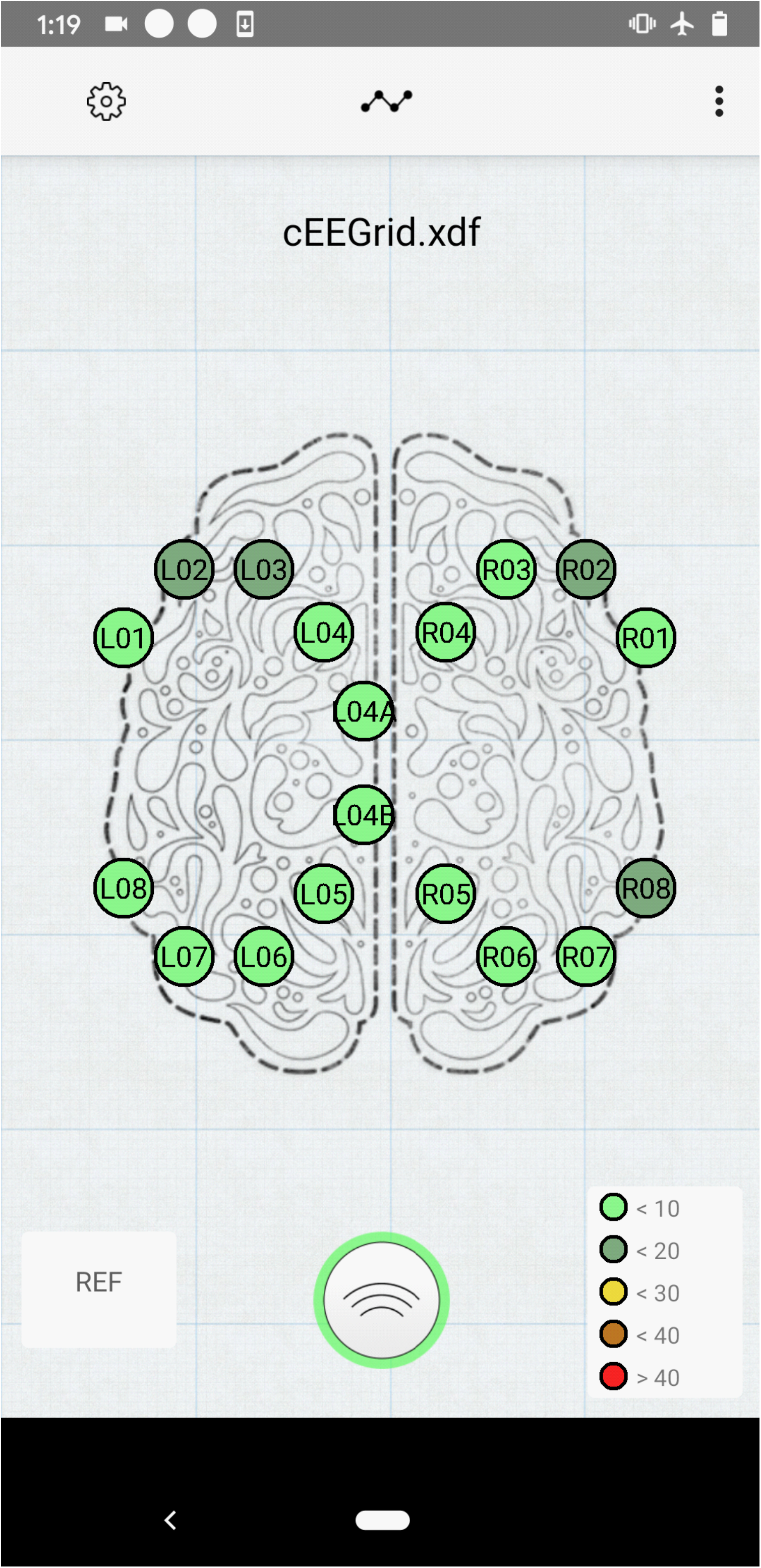
Example of good impedance. Frequently, impedance value approaches this scenario after a few minutes after placing the cEEGrid. All values are in kΩ.

**2.**
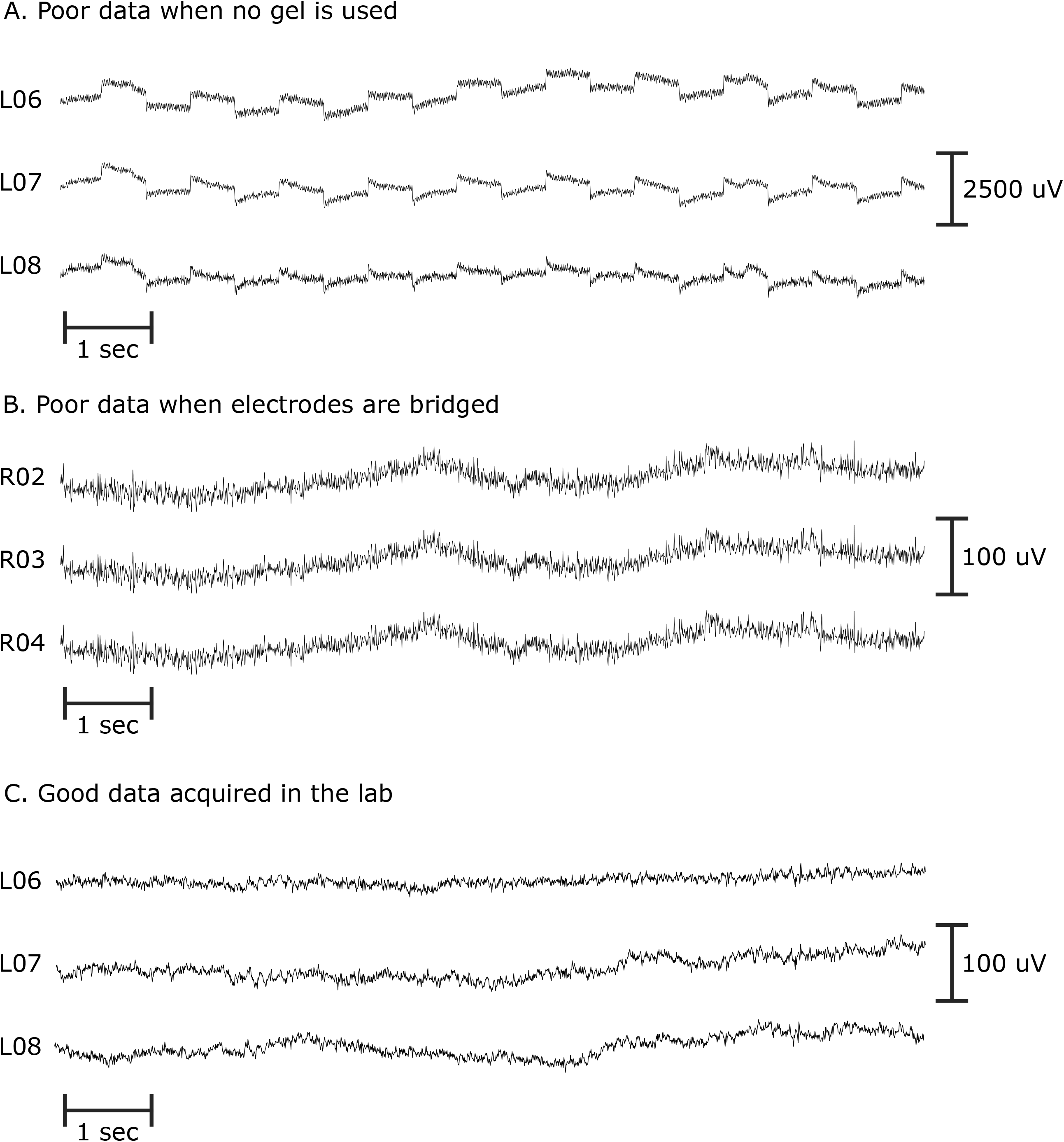
Illustration of unprocessed signals with different quality. **A**. Example of 10 seconds of data when no electrode gel is used at all. **B**. Example of 10 seconds of data when electrodes are bridged. **C**. Example of 10 seconds of good data acquired in the lab.

**3.**
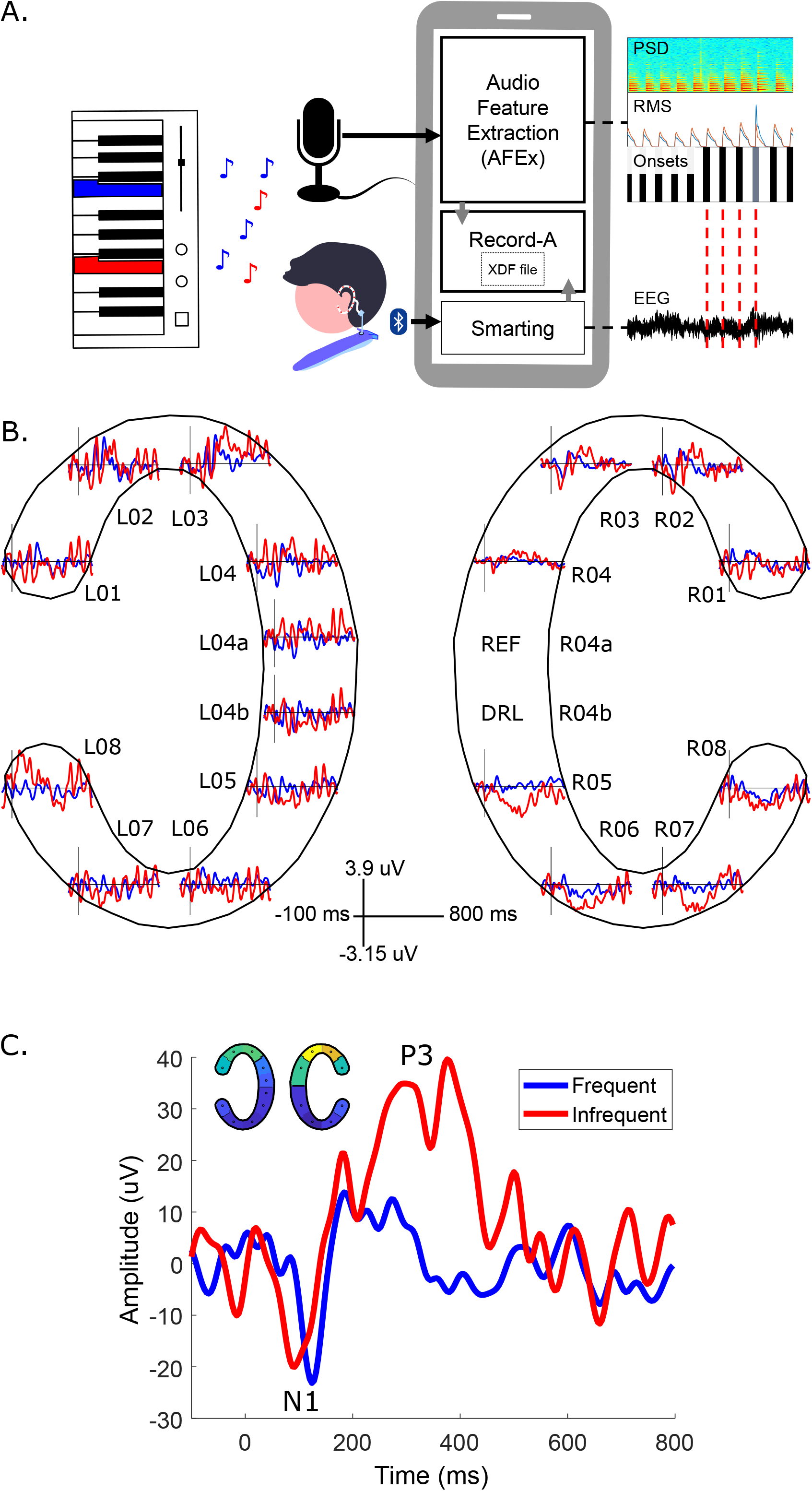
Results from an event-related potentials (ERPs) paradigm (oddball task) of one participant. **A**. The experimenter played a predefined mixture of two tones on the piano. One tone was played frequently (328 times) and another tone infrequently (78 times); the participant had to count infrequent tones. With the AFEx app, we recorded the tones, relate them to the EEG, and differentiate between infrequent and frequent tones based on the power spectral density (PSD; see ^21^ for details). The Record-A app allows to concurrently record audio and EEG^4^. The frequent tone is displayed in blue in the infrequent tone in red. The data was high-pass filtered at 0.1 Hz and low-pass filtered at 25 Hz. **B**. ERPs of all cEEGrid channels. REF: reference electrode; DRL: ground electrode. **C**. ERP based on the spatial filter displayed in the top left corner. The spatial filter was gained by using generalized eigenvector decomposition^22^ that maximizes the signal of interest. Typical components of auditory processing can be observed, such as the N1 for both tones and the P3 (marker of relevance) for the infrequent tone that had to be counted.

## DISCUSSION

We provided a protocol for ear-EEG recordings with the cEEGrid. Following the steps of this protocol ensures high-quality recordings. In the following, we draw comparisons to cap-EEG, discuss the most critical steps in the protocol along with some best practice recommendations, and we detail some modifications in using the cEEGrid.

### Comparison of cEEGrids to cap-EEG

The cEEGrid allows to unobtrusively record brain activity in everyday life settings and is well suited for longer recordings. It has several advantages compared to cap-EEG. First, due to its weight, comfort, and low visibility, it barely restricts participants in their everyday activities^1^. Second, it can be worn for extended periods of time without the electrodes drying out^1, 3, 17^, since they are sealed by the adhesive stickers. Obviously, the cEEGrid covers only a fraction of the surface of cap-EEG and therefore cannot replace EEG caps for all purposes. However, whenever a lightweight, unobtrusive, quick-to-setup solution is necessary that is minimally constricting (e.g., in the workplace), cEEGrids can provide the relevant neural information.

### Most critical steps and troubleshooting

EEG in general, and especially mobile ear-centered EEG, remains a challenging technology. Therefore, the careful preparation of the participant and placement of the cEEGrid is necessary to ensure good data quality over time. The preparation starts with the hair and the skin of the participants. The hair and skin around the ear should be washed and dried. In addition to that, the experimenter needs to carefully clean the area around the ear with abrasive gel and alcohol and ensure that the grids are firmly attached with the adhesive stickers. These steps are important and should be performed carefully to ensure a good electrode-skin adhesion and low impedances for longer periods.

Even with proper care, however, impedance for individual electrodes may be still be poor after placement of the electrodes. In general, the electrode-skin interface stabilizes over time, and we often observe that impedance decreases within 5 to 15 minutes. However, if the electrodes do not properly touch the skin (e.g. due to too much hair under an electrodes), we recommend to completely remove the cEEGrid and place it again. We do not recommend adding electrode gel to individual electrodes as this can compromise the adhesion strength of the stickers and can even lead to bridging of neighboring electrodes.

After the cEEGrid has been placed and impedance of the electrodes is low, data recording can begin. For longer recordings (> 1h), we recommend to conduct a brief data quality check at the beginning. We use, for instance, a 3-minute auditory oddball task, that can be conducted and analyzed quickly to ensure a good signal quality.

### Modifications of the method

The cEEGrid is one-size-fits-all. However, it allows for some flexibility regarding its size. By cutting the plastic of the inner side of the cEEGrid, the size of the cEEGrid may be reduced to fit larger ears. Pay special attention that you neither cut into the electrodes nor the conductive path.

Depending on the used amplifier and the recording scenario, there are different ways to place the amplifier. The fixed length of the tail of the cEEGrid and the fact that it points horizontally away from the ear limits the location where the connector to the amplifier can be placed. Different manufacturers provide adapter cables, connecting the cEEGrid to the specific amplifier (either mobile or lab-based). Different solutions have been proposed to place the amplifier; some use a headband^3^, others integrate it into a basecape^23^. For shorter experiment, a headband is suitable. For longer experiment, the amplifier can be taped to the cloths^17^ or body^2^, stored in custom-made straps, or taped to headphones worn around the neck^1^ or a neck protector commonly used for mountain biking. We have developed a prototype that combines a neckspeaker (for presenting auditory stimuli) with a mobile EEG amplifier and connectors to the cEEGrid^5^. We have used this approach successfully in a recent study (in preparation) in which we recorded ear-EEG for four hours while participants work in the office.

## Conclusion

We use the cEEGrid to investigate sound processing in everyday life^1^. Apart from this use-case, cEEGrids are especially promising for long-term recordings, for example, for diagnostic purposes such as long-term monitoring of epileptic seizures^2^, sleep staging^17^, or for attention capturing in hearing devices^8, 12^. Regardless of the desired use case, this protocol helps researchers to obtain the best possible recordings with cEEGrids.

## ACKNOWLEDGMENTS: (<u>Instructions</u>)

This work was funded by the Deutsche Forschungsgemeinschaft (DFG, German Research Foundation) under the Emmy-Noether program - BL 1591/1-1 - Project ID 411333557. We thank Suong Nguyen and Maria Stollmann for their assistance in filming the video. We thank Joanna Scanlon for the voice over in the video.

## DISCLOSURES: (<u>Instructions</u>)

The authors report no conflict of interest.

